# Convergent transcriptomic and genomic adaptation in xeric rodents

**DOI:** 10.1101/2024.10.02.616319

**Authors:** Chalopin Domitille, Rey Carine, Ganofsky Jeremy, Blin Juliana, Chevret Pascale, Mouginot Marion, Boussau Bastien, Pantalacci Sophie, Sémon Marie

**Affiliations:** LBMC, Ecole Normale Supérieure de Lyon, Université de Lyon, Lyon; IBGC, Université de Bordeaux, Bordeaux; CIRI, Ecole Normale Supérieure de Lyon, Université de Lyon, Lyon; LBBE, Université de Lyon, Lyon

## Abstract

Repeated adaptations rely in part on convergent genetic changes. The extent of convergent changes at the genomic scale is debated and may depend on the interplay between different factors. Rodents have repeatedly adapted to life in arid conditions, notably with altered renal morphology and physiology. This occurred at different time periods, allowing us to test the importance of time in convergent genomic evolution. We analyzed kidney transcriptomes from 34 species to quantify and characterize convergent evolution at the level of gene expression, tissue composition, and coding sequences. We found that several genes showed convergent expression changes, some of which also carried convergent changes in their coding sequence. We then subdivided these data to test the influence of evolutionary history. First, within the subfamily Murinae, we found more convergent gene expression, reflecting convergent changes in cell proportions. Second, we compared data for recent (within genera) and ancient (between genera) adaptations, and observed more convergent changes in the latter. Our study shows that adaptation to xeric environments in rodents involves repeated changes in tissue composition, gene expression and coding sequences, and that the degree of convergent evolution increases with both the age of the adaptations and species relatedness.

## INTRODUCTION

Repeated evolution, also known as parallel or convergent evolution, occurs when different lineages evolve similar traits independently. If the same genetic changes are used by independent lineages in repeated adaptations, the genetic basis of adaptation might be predictable. Recent genomic studies have significantly advanced our understanding of this question (Chaturvedi et al. 2022; Sackton et al. 2019; Brown et al. 2019). They highlighted a large variability in the degree of genomic convergence, which may be influenced by several factors. In particular, the amount of genomic convergence could be higher between closely related species that undergo parallel phenotypic evolution because they share a common genetic background (Bohutínská and Peichel 2023). It may also increase with the age of adaptation because species that have adapted long ago may have accumulated genomic changes affecting various phenotypic traits, which could be shared with other lineages.

Repeated adaptations to arid environments have occurred in a variety of clades. These adaptations enable species to cope with temperature and seasonal unpredictability, and with challenges to food and water availability and quality (Schwimmer and Haim 2009). They have motivated a rapidly growing area of research in genomics (Rocha et al. 2021). Studies include the comparison of renal gene expression in a few species (Bittner et al. 2022; MacManes and Eisen 2014; Giorello et al. 2018; Marra et al. 2014), dehydration experiments to study the plasticity of gene expression (Blumstein and MacManes 2023; Kim and Shin 2016; Bittner et al. 2021), genomic analyses (Cheng et al. 2023; Peng et al. 2023) and population genomic analyses (Tigano et al. 2020; L Rocha et al. 2023).

Most of the studies of adaptations to arid environments have been performed in rodents. Many xeric rodent species have acquired behavioral and physiological adaptations linked to metabolism and water retention (Rocha et al. 2021), including modified kidneys capable of producing very concentrated urine (Bankir and de Rouffignac 1985). A recent study analyzed gene expression changes and genes under positive selection in 3 independent adaptations to desert life in rodents (Bittner et al. 2022). They discovered many idiosyncratic changes but also shared changes in genes of interest known to be involved in osmoregulation and kidney function. Overall, genes involved in fat metabolism, response to insulin signaling and diabetes, stress response, endocrine system, arachidonic acid metabolic pathway and water transport have all been found to be involved in the adaptation of rodents to arid environments (Giorello et al. 2018; Bittner et al. 2022).

Here we study the repeated adaptation of rodents to life in xeric environments using transcriptomic data. We investigated the evolution of gene sequences and expression levels in kidney transcriptomes based on a large RNA sequencing (RNA-seq) dataset spanning 34 rodent species and 2 strains, including new data for 18 of them. Contrary to previous studies (Corral-Lopez et al. 2024; Bittner et al. 2022; Cossard et al. 2022; Zancolli et al. 2022; Pankey et al. 2014; Hart et al. 2018; Gallant et al. 2014; Foster et al. 2022; Parker et al. 2019b, 2019a), this dataset encompasses divergences ranging from several thousands of years to 70 million years, which provides a comparative framework for studying the effects of time scales on repeated transcriptomic and genomic evolution.

We selected 8 rodent families and a balanced number of species with xeric and mesic habitat, which allowed us to robustly infer evolutionary changes in kidney gene expression and coding sequences. First, we found that several genes carried convergent expression changes, some of which also carried convergent changes in their coding sequence. Second, we searched for genes showing convergent evolution of expression in the *Murinae* subfamily and showed that there were many more of them than in the total dataset, and that they reflected convergent changes in the proportions of renal cell types. Finally, we compared two subsampled datasets, designed to represent recent (within genera) or ancient (between genera) habitat transitions, and observed more convergent changes in ancient transitions.

## RESULTS

### Sequencing, Assembly, and Annotation

In order to investigate the evolution of kidney transcriptomes in rodents, we selected representative species belonging to 8 rodent families that diverged up to 70 million years ago (MYA, Fig. 1a, Supplemental Fig. 1). We sampled and sequenced bulk kidney RNA-seq data from 16 species and two mouse strains. In total, we generated 42 RNA-seq samples, which we combined with carefully selected publicly available RNA-seq data into a dataset of 102 samples in 34 species plus 2 strains, including samples for transcriptome assemblies and replicates for expression analyses (Supplemental Tables 1,2). Because we depend on wildlife capture, for eight species we could only secure one individual. But in most cases, closely related species from the same genus can serve as biological pseudo-replicates for the considered environmental transition. In addition, we retrieved the coding sequences from the published genomes of 24 species, obtaining in total 51 species for the coding sequence analyses, plus two strains.

**Figure 1.**
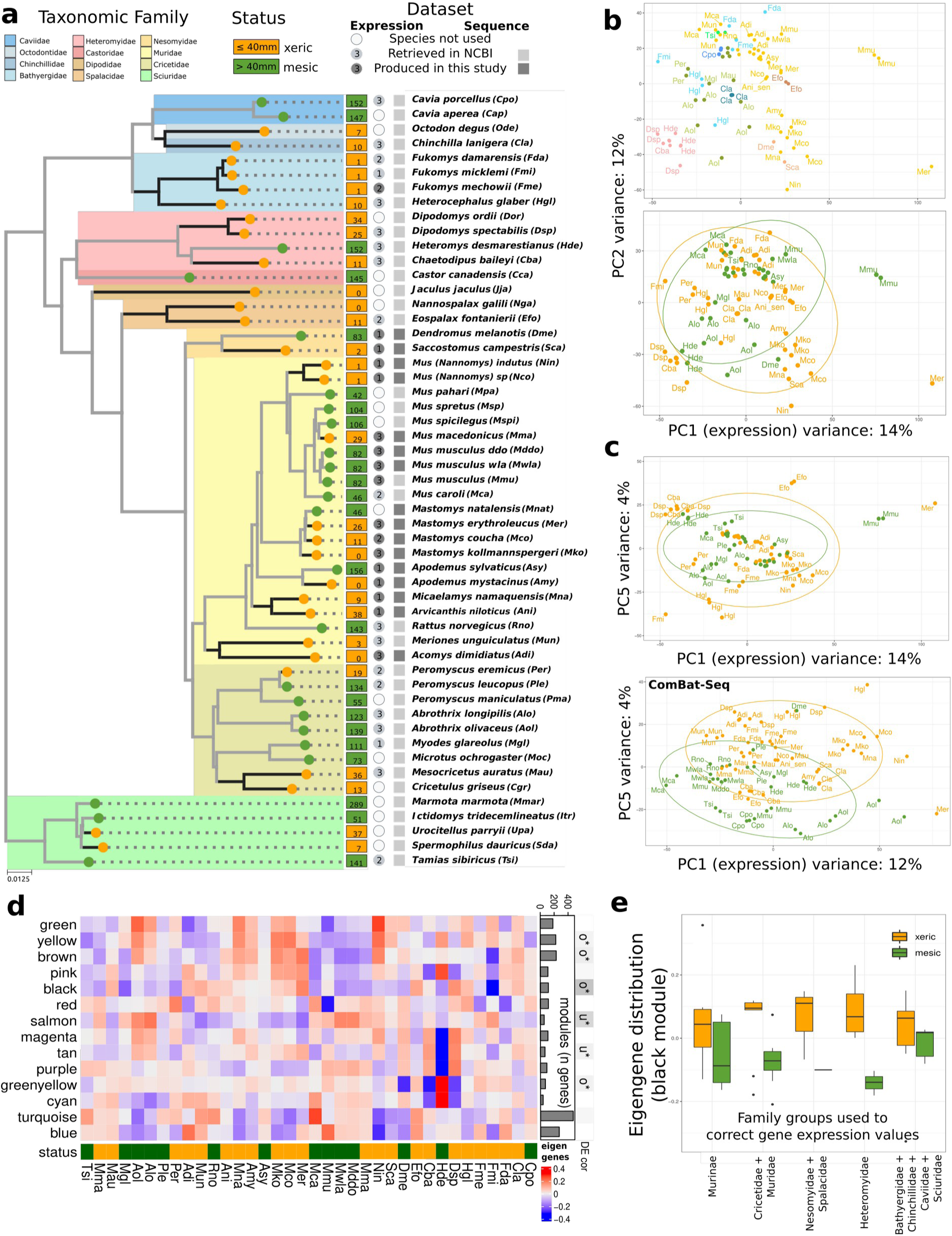
Detection of transcriptomic convergence across rodent phylogeny a) Phylogeny of the 51 species used in the study. The medians of the precipitation of the driest quarter of the year are indicated in squares defining the biological status of the species (mesic <40, green; xeric >40, orange). The number of individuals used for the expression analyses is indicated in the circles. Colors of squares and circles correspond to previously published data (light gray) and to new data (dark gray). b) First and second components of a PCA analysis using normalized but non-corrected expression values. Individuals are colored by rodent families (upper) or by habitat status (lower). c) First and fifth components of a PCA analysis using normalized and either non-corrected (upper) or batch corrected (lower) expression values. Individuals are colored by their habitat status. d) Heatmap representing eigengenes per WGCNA module. Number of genes for each module is indicated as a barplot. Modules significantly Over- (o) or Under (u)-regulated modules in xeric species are depicted at the top of the figure. e) Barplot showing distribution of eigengene values of the black module in five phylogenetic groups.

After constructing *de novo* transcriptome assemblies of the RNA-seq data and assessing their quality (Supplemental Table 3, 4), we derived gene orthology relationships between all species and isolated 11437 gene orthogroups with at least 3 species. We performed gene expression analyses based on these 1:1 orthologs and on the high quality RNA-seq dataset (80 samples) and reconstructed gene alignments and phylogenetic trees for coding sequence analyses (see Methods, Supplemental Fig. 2).

To associate each species with a biological status corresponding to xeric and mesic life, we determined its geographical distribution area and extracted the corresponding bioclimatic variables. Because an annual average pluviometry can hide large differences between seasons, we decided to use the precipitation of the driest quarter of the year and to define a species with less than 40 mm as a xeric species. We annotated the status of 1898 species along a published rodent phylogenetic tree containing 2260 rodent species (Fabre et al. 2012) and modeled state transitions to infer ancestral states at each internal node of the phylogeny. We then extracted the ancestral states for the subset of nodes corresponding to our dataset (see Methods, Supplemental Fig. 3,4). We annotated 29 and 22 xeric species in the coding sequences and expression datasets, respectively (Fig. 1a).

### Characterization of global patterns of convergent expression in rodent kidneys

The first components of a principal component analysis (PCA) on expression levels tended to group samples from the same species together and to separate samples from different rodent families (Fig. 1b). The difference between species accounted for 89.7% of the total variation, and the difference between families for 37.0% (between class analyses). This suggested that gene expression diverged following the phylogeny, as seen previously in several studies including rodents (Bittner et al. 2022).

To minimize the influence of phylogenetic effect we applied a batch correction using *ComBat_seq* to account for the effect of the phylogeny at the family level (see Methods). We performed another PCA with these phylogeny-corrected data. We observed that the fifth principal component, accounting for 4% of the variance, separated xeric and mesic species, although incompletely (Fig. 1c). This shows that some gene expression levels in xeric species have converged and acquired similarities.

We quantified differential expression by using pairwise contrast between xeric and mesic status using these phylogeny-corrected data and found 26 genes significant at the 0.1 adjusted p-value threshold and with log-fold change greater than 0.4 (21 genes were found with LFC>1, Supplemental Table 6). To assess whether this number of genes was larger than expected by chance, we compared it to the numbers measured in 1,000 permuted datasets (permutation method adapted from (Bittner et al. 2022), see Supplementary Methods). We found that the true number of differentially expressed genes is eleven times higher than expected (p-value < 0.0001). Unsurprisingly given its modest size, this group of 26 genes revealed only two overrepresented Gene Ontology (GO BP) terms, “small molecule biosynthetic process” and “animal organ morphogenesis” (adjusted p-value < 0.02). Among these genes, we found two members of the solute carrier (SLC) gene family, Slc35b4 and Slc40a1 (Kordonowy and MacManes 2017), which is marginally more than expected by chance (Fisher exact test, p-value = 0.059). Slc40a1 is an iron exporter previously identified in a dehydration experiment (Kordonowy and MacManes 2017). This set also included 5 genes known as kidney markers or associated with renal diseases, Casr (associated with hypocalcemia and calcium kidney stones (Vezzoli et al. 2011; Hanna et al. 2021)), Ctsh, Xpnpep2 (Böttinger 2010), Fam20a (associated with enamel renal syndrome (Wang et al. 2014)) and Cpne2 (renal cancer (Zhou et al. 2018)).

We hypothesized convergence in gene expression could be detected in functionally related modules of genes, which work together in the kidney and therefore may tend to change their expression in a coordinated manner along the phylogeny. We ran a correlation network analysis (Langfelder and Horvath 2012) on the complete expression dataset and found 14 modules of co-varying genes. In the following, these modules are given arbitrary color names and represented by their eigengenes, which correspond to the weighted mean of expression levels in the module. Six of them correlated significantly with the aridity status (Fig. 1d). Within each rodent family, the expression level in modules is distinct between xeric and mesic species, confirming that a convergence signal is present alongside the phylogenetic signal. We looked for functional overrepresentation for GO terms and reactome pathways within the modules. For example, the black module (Fig. 1e) was related to blood vessel development and extracellular matrix organization, the salmon module was related to metabolic processes and the green-yellow module to metabolic processes and ion homeostasis (Supplemental Fig. 5 and Supplemental Table 7).

### Global patterns of convergent evolution in coding sequences

To characterize cases of convergent evolution in coding sequences, we searched for sites where preferred amino acids differ between mesic and xeric species. We used Pelican, a method that takes into account the phylogeny of the species and which proved to be the best of its kind in a recent benchmark (Duchemin et al. 2023). We selected 4,065 gene families with well-aligned single-copy orthologs found in at least 20 mesic and 25 xeric species. We further refined the list of candidate sites by discarding all sites that had undergone a substitution in only one of the xeric clades, considering that we were interested in profile changes that have occurred in a convergent manner, at least in two xeric clades. We then ranked the genes based on the best p-value among their sites and studied their functional relevance by using gene set enrichment analyses (GSEA). Three pathways of the Reactome database displayed a significant enrichment: SLC-mediated transmembrane transport, fatty acid metabolism and transport of small molecules (adjusted p-value < 0.1, Supplemental Fig. 6 and 7 for corresponding enriched GO and REACTOME terms). The genes with the best detected sites were also enriched for the set of differentially expressed genes (GSEA, p-value = 0.0269, Supplemental Fig. 8). This enrichment was supported by 12 genes whose gene expression levels and amino-acid profiles differed between xeric and mesic species (Fig. 2). However, repeated evolutions were not observed in all independent transitions, but limited to a subset of the families of rodents. This suggested that convergent evolution might be more important when examined within a family.

**Figure 2.**
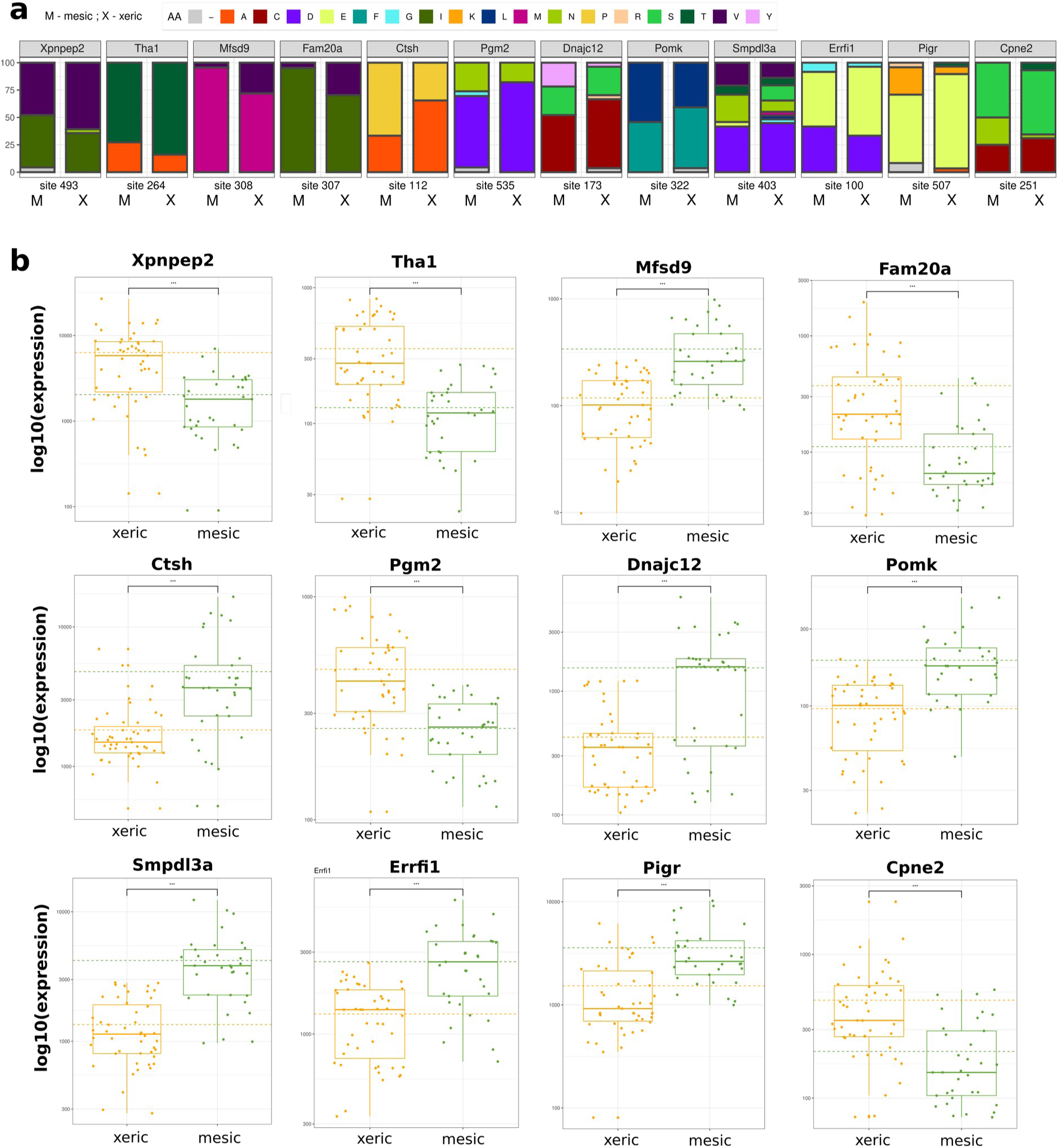
Twelve core genes with convergence detected in the coding sequence and in expression. a) Amino acid composition (percentage) for the best site of the twelve genes in function of mesic or xeric species. b) Boxplots showing normalized expression of the twelve genes. Orange dashed line shows the mean expression of xeric individuals and green dashed line shows the mean expression of mesic individuals.

### Convergent evolution in gene expression, coding sequences, and cell proportions in the Murinae subfamily

We investigated convergent evolution within the Murinae, a large subfamily of rodents that diversified quickly after it originated 11.2 MYA (Aghová et al. 2018). This left us with a dataset of 7735 genes and 14 taxa including 8 xeric species to study gene expression, representing 4 independent habitat transitions (see Methods, Fig. 3a). Because this dataset contains species for a single rodent family, we did not apply our family-level phylogenetic correction. The first two components of the principal component analysis of this dataset showed a clear distinction between mesic and xeric species (Fig. 3b). Three individuals of the xeric species *Mus macedonicus* locate with mesic species of the genus *Mus*, probably because they are closely related and because *M. macedonicus* lives in the moderately xeric mediterranean environment. Apart from this, the first component carries most of the separation between xeric and mesic individuals and accounts for a large proportion of the variation (24%). This suggests that there is a conserved and pervasive habitat-related transcriptomic signature that rivals phylogenetic divergence, within Murinae.

**Figure 3.**
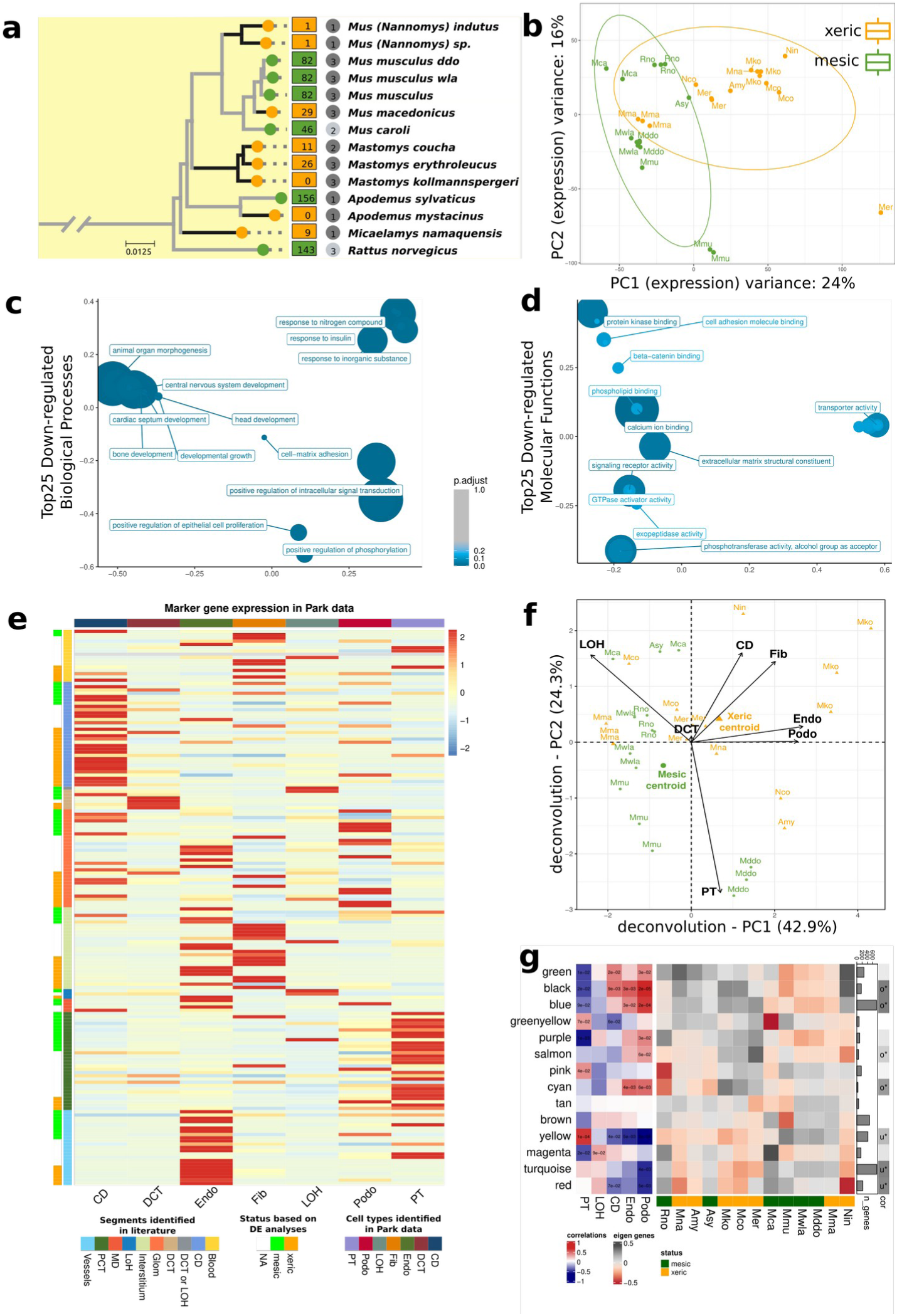
Detection of transcriptomic convergence in the Murinae subfamily. a) Phylogenetic relationships of the Murinae species and strains used. The biological status of the species is indicated as in fig. 1. b) Visualization of the two first components of the Principal Component Analysis (PCA). c-d) Clustering of the top 25 overrepresented GO annotations for biological processes (c) and molecular functions (d) using genes down-regulated in xeric species. e) Heatmap showing expression of cell type specific markers retrieved from the literature (rows) in the single-cell dataset of Park et al. (column). Environmental status is indicated if the marker genes are found in differential expression analyses. f) PCA summarizing cell proportions estimated by deconvolution. Centroid values from both xeric and mesic are indicated. Cell types included collecting duct (CD), proximal tubule (PT), loop of Henle (LOH), distal convoluted tubules (DCT), podocytes (Podo), endothelial cells (Endo, that also contain descending loop of Henle) and Fibroblasts (Fib). g) Heatmap representing eigengenes per WGCNA module. Number of genes for each module is indicated as a barplot. Significant Over- (o) or Down (d)-regulated modules are depicted at the right side of the figure. The left heatmap represents Pearson correlations between values of the deconvoluted proportions and module eigengenes.

We quantified differential expression between xeric and mesic species and obtained 692 genes with significant differences, which is 19.8-fold more genes than expected and highly significant (p-value < 0.0001, Supplemental Table 6).

We intersected this list with marker genes of kidney cell types and genes associated with renal diseases. We found 30 marker genes and 20 disease genes in our list, 1.4-2 times more than expected (p-values 0.0003 and 0.17 respectively, see Methods and Supplemental Table 8). Differentially expressed genes included 2 aquaporins (Aqp2, a vasopressin-regulated water channel involved in diseases affecting urine-concentrating ability (Pannabecker 2015) and Aqp7, expressed in proximal tubules, with phenotypes of insulin resistance and important in glycerol reabsorption in the kidney (Sohara et al. 2006)), 18 solute carriers (including the urea transporter Slc14a2, the sugar transporter Slc17a5, the sodium carrier gene Slc8b1, Slc27a2 that plays an important role in hepatic fatty acid uptake and was found overexpressed in kangaroo rat kidney (Marra et al. 2012)), and 6 genes of the arachidonic acid pathway. Focusing on genes significantly down-regulated in xeric species, enriched GO terms included response to insulin, regulation of autophagy, transmembrane transporters (Fig. 3b,d).

We performed a correlation analysis and found 14 coexpression modules in the dataset, 7 of which significantly correlated to xeric/mesic status, even though a phylogenetic effect was also visible (see for instance the Mus clade, Fig. 3g). The 7 modules presented functional categories congruent with the differentially expressed genes, such as glucose metabolism (green-yellow module), regulation of apoptotic processes and response to lipids (blue, Fig. 3h), solute carriers (yellow) (complete lists are available in Supplemental Fig. 5).

Bulk RNA-seq data reflects variation both in expression per cell and in cell type composition. Here, different species may exhibit divergent tissue histologies as part of their adaptation to the xeric environment. We therefore decided to deconvolve the bulk RNA-seq data to investigate changes in cell type composition distinguishing xeric and mesic species (see methods and Supplemental Fig. 9). Cell proportions were estimated by MuSiC (Wang et al. 2019) using published kidney single-cell RNA-seq data from mouse (Park et al. 2018). The first two components of a PCA calculated on these proportions separated xeric and mesic species (Fig. 3f), with the exception of *Mus macedonicus* samples (which resemble mesic species as already seen above), one of the *Mus caroli* samples and our single sample of *Apodemus sylvaticus*. Cell types that mostly contributed to this axis were, on the xeric side, collecting duct cells (CD), podocytes (Podo) and endothelial cells (which also contain LOH cells) and on the mesic side, proximal tubule cells (PT). This discrimination is significant (discriminant analysis, p-value = 0.002, Monte-Carlo test based on 1000 replicates) and consistent with biases observed between xeric and mesic species in the expression of 177 marker genes (Fig. 3E and Supplemental Table 8).

We characterized sites with convergent evolution in coding sequences on 3670 gene families with more than 7 xeric and 9 mesic species and studied their functional enrichments by using GSEA. Three pathways of the Reactome database displayed a significant enrichment: SLC-mediated transmembrane transport, fatty acid metabolism, and transport of small molecules (adjusted p-value < 0.1, Supplemental Fig. 6). Differentially expressed genes displayed a marginal enrichment (p-value = 0.051, Supplemental Fig. 8).

Much more genomic convergence was observed at the level of a single rodent family than in the entire dataset. One possible reason is that Murinae have recently diverged and thus share a common genomic background. Another reason may be that their adaptations all occurred in a similar time frame, whereas in the entire dataset some species belong to lineages that have adapted over tens of millions of years to extreme environments, while others adapted very recently from mesic ancestors to moderately xeric environments.

### Comparing datasets with ancient (between genera) and recent (within genera) adaptations

We prepared two datasets of similar size to that of Murinae, in terms of number of species and number of transitions, but where these transitions to xeric habitat are either relatively recent (within the same genus and younger than 6 MYA), or more ancient (at the base of a rodent family and/or older than 6 MYA, see Supplemental Fig. 1).

The subset with “within-genera” transitions allowed us to study gene expression levels in 15 species, representing four recent transitions to the xeric condition in two sister families (Fig. 4a). The fourth PCA axis correlated best, although imperfectly, with xeric/mesic status and accounted for 9% of the variation (Fig. 4c). We found only 29 genes showing evidence of differential expression, which nevertheless constituted a significant enrichment (11-fold, p < 0.0001). Co-expression analyses identified 15 modules, of which only one module was significantly correlated with aridity state (Fig. 4d). 29 species were available for analyzing convergent sequence evolution, spanning 4 transitions (4604 gene families, with at least 20 mesic and 4 xeric species). The sites we detected were not enriched in differentially expressed genes (p-value = 0.12), nor in the module of coexpressed genes correlated with aridity status (Supplemental Fig. 8).

**Figure 4.**
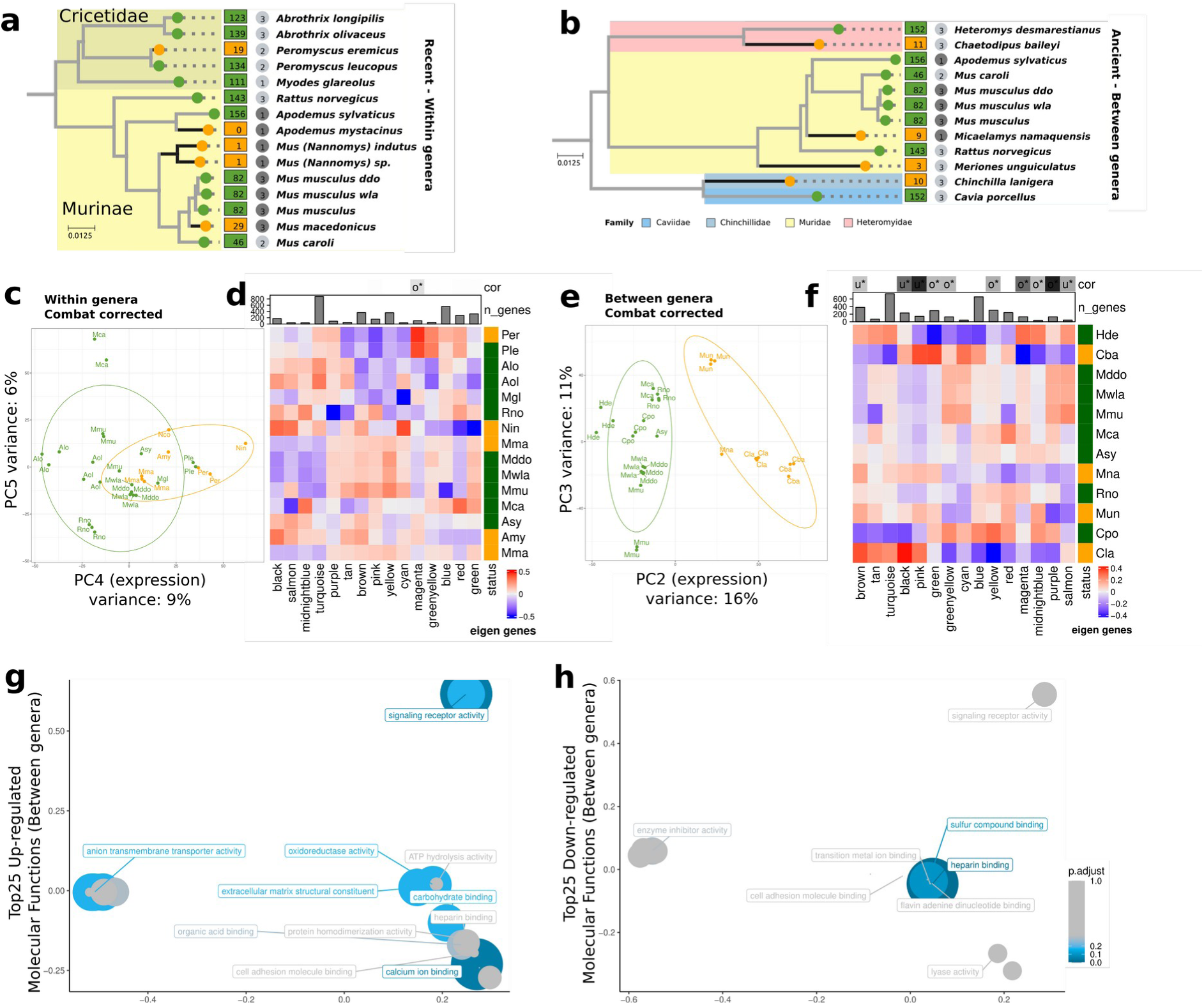
Detection of convergence in datasets with ancient (between genera) and recent (within genera) transitions. a and b) Phylogenetic relationships of the sets containing recent (within genera) and ancient (between genera) transitions, respectively. The biological status of the species is indicated as in fig. 1. c and e) PCA plot using corrected expression values from the set of recent and ancient transitions respectively. d and f) Heatmap representing eigengenes per species and per WGCNA module in the set with recent and ancient transitions respectively. g-h) Clustering of the top 25 overrepresented GO annotations for molecular functions and associated with genes up (g) and down (h)-regulated in xeric species in the dataset with ancient transitions.

The subset with “between-genera” transitions contained 12 species belonging to 4 families, and representing 4 ancient xeric transitions (Fig. 4b). The second axis of the PCA clearly separated xeric and mesic species (16% of the variation, Fig. 4D). There were 632 differentially expressed genes (3-fold excess based on permutation test, p<0.0001). Up-regulated genes were involved in anion transporter activity, oxidoreductase activity, organic acid and calcium binding (Fig. 4g). We identified 15 modules of co-expressed genes, of which 10 are correlated with the aridity state, with concordant functional enrichment (Fig. 4f, such as regulation of glucose metabolic process for salmon, solute-carrier-mediated transmembrane transport for green-yellow, see Supplemental Table 7). 28 species were available for analyzing convergent sequence evolution (3670 gene families with at least 20 mesic and 4 xeric species). The sites we detected were enriched for the set of differentially expressed genes (p-value = 0.033) and for three modules of coexpression which are all correlated with aridity state (Supplemental Fig. 8).

For certain families important in renal function (ABC transporters, aquaporins, ion transport, solute carriers), we examined the 5 best sites per gene. We retained 9 genes (Fig. 5), either because the amino-acid profile changed markedly in one position for 2 to 3 xeric transitions, or (most often) because several positions behaved in a correlated manner, suggesting the structure of the protein might have changed. The gene Ctns (a lysosomal transporter causing kidney failure characterized by proximal tubular dysfunction (Attard et al. 1999)) displays a convergent change at 3 sites in 2 branches.The same change occurred in two other xeric transitions in the total dataset (Supplemental Fig. 10). This gene is not differentially expressed in the “ancient transition” (between genera) dataset but gene expression is significantly upregulated in Xeric species in the “murinae” dataset. The gene Abcc2 displayed a site with convergent evolution in three xeric species. In humans, mutations in this gene are associated with substrate transport efficiency in the kidney (Muhrez et al. 2017). Slc28a1 (nucleoside transport in kidney (Persaud et al. 2023)) and Slc36a1 (aminoacid reabsorption in proximal tubule (Chrysopoulou and Rinschen 2024), significantly down-regulated in xeric species in the Murinae dataset) both displayed convergent evolution in two lineages. Scnn1a (mutations in this gene impact sodium balance in mouse and human (Rossier et al. 2002)) is an example where the same amino-acid tends to increase in frequency at different positions of the protein. For Slc2a4, we observed a well-conserved sequence among mesic species, but more variation in xeric species, suggesting a relaxation of selection. Therefore, many different patterns of convergent evolution in amino-acid profiles are present in the data.

**Figure 5.**
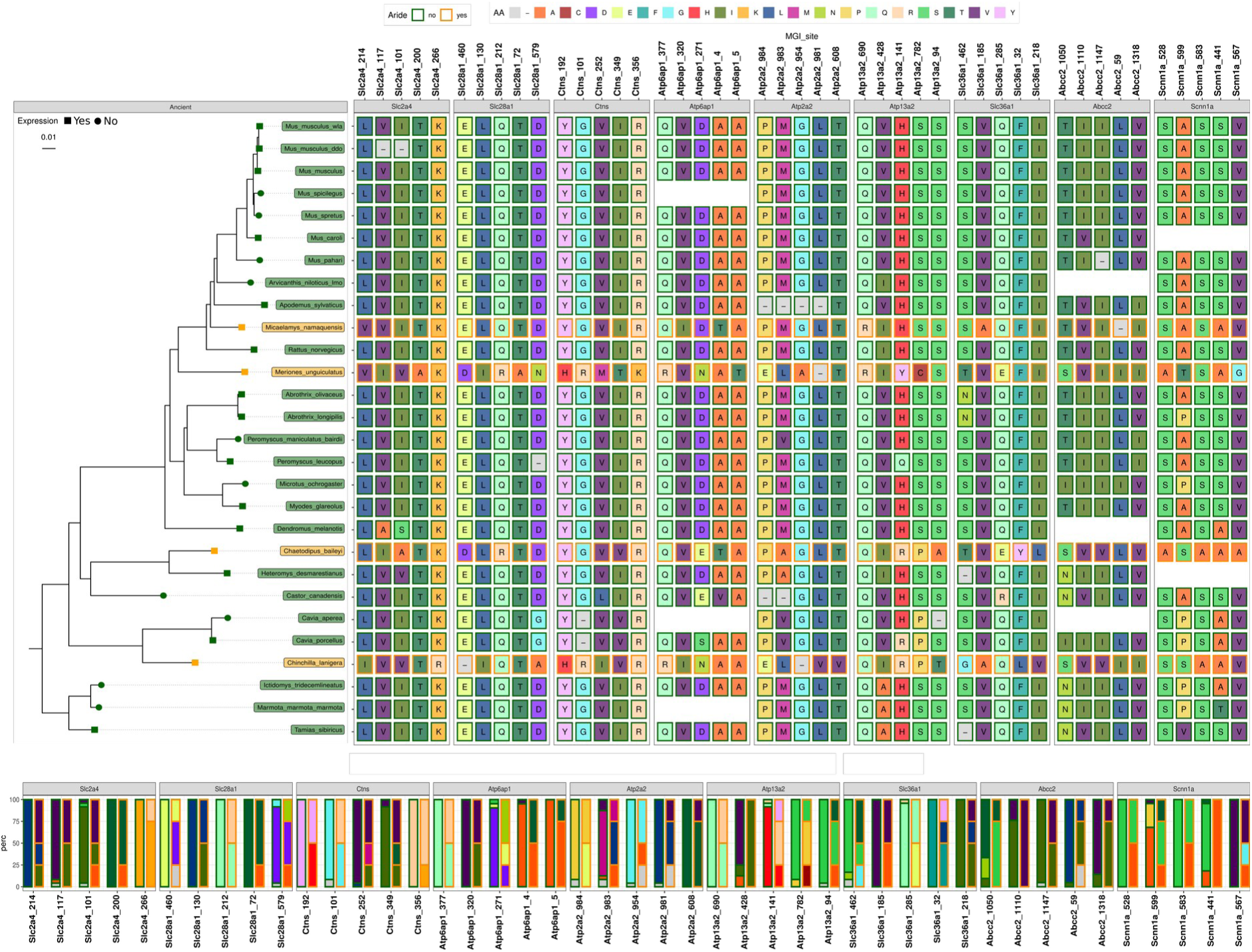
Selected sites with evidence of convergent shift in amino-acid profiles detected by Pelican. (Top) The 5 best sites are represented for 9 genes from relevant gene families. Amino-acids are represented by squares of different colors. Blank spaces indicate that the sequence was missing for that species. Amino-acids from xeric and mesic species are circled in orange and green, respectively. (Bottom) Amino acid composition (percentage) for each site in function of mesic or xeric species.

### Comparison of differentially expressed genes between datasets

We compared the genes with repeated changes in expression in different datasets to see whether the processes involved are the same. There were only 4 common genes between our 4 datasets: Two genes up-regulated in xeric species, Cpne2 and Fam20a, and two genes down-regulated in xeric species, Ctsh, Casr (a marker of the distal tubules that regulates calcium reabsorption (Habuka et al. 2014; Vezzoli et al. 2019)), of which 3 show some degree of convergent amino-acid profiles in at least one site (Fig. 2).

We then focused on the datasets with most differentially expressed genes, the Murinae (692 genes) and the “ancient transitions” datasets (632 genes) and we found an overlap of 84 for differentially expressed genes, of which 71 are biased in the same sense. This represented a 2.1-fold increase and a significant enrichment (Chi-squared test, p-value<10-9), although 6 taxa are found in both datasets. There was no particular functional enrichment within ancient-specific genes. The common genes were significantly overrepresented in “transmembrane signaling receptor activity”, while the murine-specific genes were enriched in “lipid transporter activity”, “cellular response to oxygen−containing compound”, “epithelial cell apoptotic process” and “innate immune system” (complete list of GO terms in Supplemental Fig. 7).

## DISCUSSION

We studied repeated genome and transcriptome evolution in response to adaptation to aridity, at the macroevolutionary scale, in 8 families of rodents. Our analyses covered transcriptomes, coding sequences and cell type proportions. Together with our extensive species sampling we studied convergent molecular evolution at multiple levels and across time scales. The strength of our study is the large number of mesic and arid species including already available transcriptomic data as well as a wildlife sampling that captured 1-3 individuals for several species. A caveat in this strategy is that we only have a single individual in several species. We fully acknowledge that this prevents studying species-specific changes in expression, but it does not prevent concluding on convergence at the clade level. Indeed in most cases, sister species that share the same ancestral environmental transition represent biological variation in the branch and serve as pseudo-replicate to quantify convergent evolution.

### Gene functions and overlap with previous studies

Our expression comparisons revealed a significant amount of genes associated with kidney physiology. Among the common physiological systems allowing mammalian survival in deserts described in a recent survey, there were increased urine osmolarity and increased water reabsorption from the kidney, higher levels of plasma creatinine, increased plasma osmolality, change in insulin secretion for adaptive tolerance to dehydration and starvation (Rocha et al. 2021). We found in our data several genes and pathways relevant to these systems.

Aquaporins form a gene family of water transporters that has been associated with desert adaptation in rodents (Bittner et al. 2022; Pannabecker 2015; Marra et al. 2014; Giorello et al. 2018). In the Murinae dataset, we found convergent upregulation of Aqp2 and Aqp7 in xeric species. Aqp2 is the dominant water transport gene in the medullary Collecting Ducts. Since its spatial pattern of expression seems similar in many rodent species (Pannabecker 2013), we may have detected a change in intracellular expression level. Of note, because for some species we rely on *de novo* transcriptome assemblies, we cannot reconstruct the sequences of genes with very low levels of expression. Aqp4 for instance, another important water transporter (Donald and Pannabecker 2015), is not available in our datasets, possibly for this reason. In a previous study, aquaporin expressions were shown to respond to hydric stress (MacManes 2017), but in our dataset we cannot discriminate between adaptation and plastic response.

We found that many solute carriers are differentially expressed. Slc14a2, which was upregulated in xeric species in Murinae, is an urea transporter whose knock-out causes decreased urine osmolality (Fenton et al. 2004). Slc8b1, a calcium:sodium exchanger, was upregulated in xeric species in Murinae and carried marks of positive selection in a previous study of adaptation to aridity in *Peromyscus* rodents (Tigano et al. 2020).

We intersected our differentially expressed genes with results from a recent study of convergent adaptation to desert life in 3 pairs of rodent species (Bittner et al. 2022). Among their list of genes with evidence for convergent differential expression and involvement in kidney physiology and/or signature of sequence selection, we found that four genes were also differentially expressed in the Murinae dataset (Fstl1, Cpne2, Paox and Blmh).

### Convergent evolution in cell-type proportions

The structure of kidneys varies considerably among mammals (Zhou et al. 2023), with differences in renal histology related to adaptations to the xeric environment (Bankir and de Rouffignac 1985). Certain differences are species-specific, such as the unique papillary loop of the chinchilla (Chou et al. 1993). Others have been measured across a wide range of species, such as the relative thickness of the medulla, which is proportional to the maximal length of the loop of Henle (Beuchat 1996) and is positively associated with habitat aridity, once body mass and phylogenetic signal are accounted for (al-Kahtani et al. 2004). Hence, when we deconvolved our bulk kidney RNA-seq to estimate kidney cell type composition, we were expecting to observe an increased proportion of cells from the loops of Henle in xeric species. We do observe a signal of convergence in several cell types. Indeed LOH cells (actually, cells from the ascending loop of Henle) and endothelial cells (which also include LOH cells, but from the descending loop), but also collecting duct and distal collecting duct cells (CD, DCT), and Podocytes tend to be in higher proportion in xeric species. The proportion of proximal tubule cells (PT) was enriched in mesic species. The convergent changes in proportions are consistent with convergent changes in many marker genes. This is for instance the case for the internal medullary collecting duct (CD), a cell type that selectively expresses Aqp2 (Chen et al. 2017; Habuka et al. 2014; Miao et al. 2021). We found that xeric Murinae species express Aqp2 at a significantly higher level in bulk RNA-seq data and, accordingly, CD is found in a higher relative proportion. Conversely, Slc28a1, a marker gene of the proximal tubule (PT), is downregulated in xeric murinae, in accordance with a smaller number of PT cells in these species.

Beyond histological differences, this signal could also be explained by subtle differences in cell type annotation. We lack resolution in the granularity of cell type annotations, particularly between PT segments (Chrysopoulou and Rinschen 2024) and between short and long loops of Henle. In species with high capacity for urine concentration, the relative number of short loops is increased (Pannabecker 2013). Another possibility is that cell type identity has shifted along the loops. For instance, as compared to rats, Aqp1 was found to be expressed in a greater territory of the descending thin limbs of the loops of Henle in the kangaroo rats, which may allow greater solute concentration (Urity et al. 2012). In our deconvolutions, we do not have the precision necessary to test the above hypotheses.

Variation in cell proportions estimated by deconvolution is correlated with global variation in gene expression, as evidenced by PCA axes and coexpression modules. This reiterates the often overlooked impact of differences in cellular proportions on bulk RNA-seq. Renal single-cell RNA sequencing data from multiple rodent species, ideally including xeric species, will be needed to harness the full power of deconvolutions in our system. As in the kidney, cellular composition has likely undergone convergent changes in many other complex and heterogeneous organs and the deconvolution approach we present here could help study them.

### Convergence in amino-acid profiles

A few convergent phenotypes, such as echolocation or C4 carbon fixation in plants, provide classic examples of perfect convergent amino-acid sites (Besnard et al. 2009; Marcovitz et al. 2019). Since this definition is very restrictive, we wanted a method that can identify these sites as well as others with more flexible criteria. We used Pelican, a new method that relies on amino-acid profiles to identify sites that would correlate with xeric and mesic habitat along the species phylogeny (Duchemin et al. 2023). It was not possible to compare the number of sites between the different subsets because Pelican’s p-values are not calibrated, but we were able to rank the sites based on their scores. Even in the best sites, we did not observe sites with the exact same amino-acid change occurring in all xeric species. We do not think this is due to a lack of sensitivity, as suggested by simulations on trees whose depth and number of transitions are comparable to ours (Duchemin et al. 2023). Consistent with our results, a recent analysis of molecular evolution associated with diverse convergent phenotypes in rodents found very few cases of perfect convergent amino acid evolution (Roycroft et al. 2021). We were not able to study all the genes in the rodent genome, since we left aside genes that had undergone recent duplications and low-expressed genes whose sequence could not be reconstructed in certain species. It therefore remains possible that examples of perfect convergent amino acid substitutions are hidden among the remaining genes.

Gene set enrichment analyses, based on Pelican site ranking, identified pathways relevant to xeric adaptations, such as SLC-mediated transmembrane transport, fatty acid metabolism, small molecule transport and lipid metabolism. We described above a modest but significant overlap between the sites detected by Pelican and the lists of differentially expressed genes. We did not expect perfect overlap since, in theory, differentially expressed genes in the kidney correspond primarily to processes in renal physiology, while amino acid changes may relate to various aspects of the adaptation to xeric lifestyle, possibly outside of the kidney.

### Effect of time on the convergent evolution of expression levels

Several studies have now shown that cases of repeated phenotypic evolution exhibit higher rates of convergent molecular evolution within recently diverged lineages than within lineages that diverged a longer time ago, but the relationship becomes less clear within clades with older divergence (Bohutínská and Peichel 2023). Here we took advantage of the fact that we sampled many species to study the effect of time scales on convergent expression evolution at the macroevolutionary level. We focused on a single organ and a single rodent clade. Compared to a meta-analysis of different works carried out in different clades and for different phenotypic traits, this has the advantage of better controlling the confounding effects of differences in polygeny and genome architecture on the level of molecular convergence.

We studied two different time effects, the age of adaptation and the age of the most recent common ancestor from which different lineages have adapted. We studied the whole dataset, and 3 subtrees with roughly the same number of leaves, xeric species, and habitat transitions. In all 4 datasets, we observed an excess of convergent changes in gene expression as compared to expectations. However, much larger sets of shared changes in gene regulation were observed when convergent evolution was detected within a single rodent subfamily (Murinae), or when we compared relatively old adaptations to xeric lifestyle (between genera) to relatively recent adaptations (within genera).

The reasons for this excess may differ in the two cases. The convergences that have taken place within the Murinae subfamily are perhaps favored by the fact that these species share a relatively recent common ancestor and therefore still have a similar genomic background. This results in similarities in their mutational landscape, protein interactomes, regulatory pathways, or even in some cases in shared alleles. Thus, adaptive changes are more likely in certain genes because the genetic structure, or the probability of specific mutations, is more favorable to them (Schluter 1996). A phylogenetic effect is visible in the coexpression modules, which can be considered as the mark of this common background in gene expression.

The molecular convergences between distant lineages that have long adapted to aridity could be attributed to the fact that many important changes in physiology have accumulated, increasing the chances of finding some repeatedly. We observed a significant overlap between convergent genes in the “between-genera” and Murinae datasets, but also murinae-specific functional enrichments. This highlights that the amount of convergent evolution is influenced by both historical contingency, leading to clade-specific adaptations, and time scale.

## METHODS

### Wild and lab maintained sampling

To cover a maximum of rodent families, we performed a sampling of wild and lab maintained rodent species. The collected individuals are summarized in Table S2. We were able to retrieve RNA-later preserved kidneys (see next section) from three different laboratories and from several natural habitats over six countries (Senegal, South Africa, Cameroun, Nigeria, Benin, Greece, France). Because many domestic mouse samples exist in the databases, we generated our own samples. Moreover, we chose to add two strains of *Mus musculus*, namely DDO and WLA, that were maintained for generations at the “Conservatoire de la souris” (Montpellier, France). The DDO strain was initially captured in Odis (Denmark). The WLA strain, initially captured in Toulouse (France). We selected the different Mus species from the “Conservatoire de la souris” to obtain a range of consumption. Interestingly, we obtained two genera, i.e. *Mus* and *Mastomys*, from the Murinae family with at least four different species.

### Kidney dissection

To homogenize dissections between the different collectors, we set up a specific protocol. The main objective was to avoid introducing any bias in gene expression by recovering RNA from subparts of the kidney that would not be representative of the whole organ, or by co-preparing other tissues, such as adrenal gland or fat, with the kidney. Animals were mostly captured during the night or early morning and killed using cervical dislocation for small animals and a lethal intracardiac dose of pentobarbital for bigger animals administered under deep anesthesia. Immediately after, the kidneys were dissected. Adrenal glands were carefully removed as well as fat using a stereomicroscope when available. Dissections were carried out in a petri dish placed on ice, with cold cell culture medium, or PBS or HBSS solution. Kidneys were then transferred in a small cell culture dish with RNA later (THERMOFISHER – AMBION solution, AM7020) and cut in small pieces of approximately 2-3mm^3^. The pieces with the RNA later were then transferred to 2 mL (or 14 mL depending on the size of the kidney) tubes with at least 5-10 volumes of RNA later. When possible, tubes were agitated overnight at 4°C on a rocker and then stored at −20°C. For field captures, samples were occasionally kept at 4°C for 1-2 days.

### RNA extraction and sequencing

We prepared RNA-seq libraries for 42 samples corresponding to 16 rodent species and 2 mouse strains (Table S1, S2, S4). For representativity, we used the whole kidney, including for large-sized species. All pieces from a single kidney were lysed in trizol with a Precellys homogenizer (Bertin). When needed, several lysates were prepared independently, and then carefully mixed together to ensure homogeneity of the lysate before precipitation and further purification using the RNeasy mini kit from QIAGEN. RNA integrity was controlled on a Tapestation (Agilent Technologies, most samples had a RIN between 7.8-10, a few samples had a RIN between 6.5 and 7.1 RIN over 6.5 were selected). PolyA+ libraries of the large-scale dataset were prepared with the Truseq V2 kit (Illumina, non stranded protocol), starting with 150 ng total RNA. Libraries were sequenced (Illumina HiSeq4000, 100bp paired-end or 50bp single-end reads, see Table S2). We evenly distributed 10 samples on 5 lanes for single-end libraries and 6 samples on 4 lanes for paired-end libraries.

### Bioclimatic variables

We obtained the geographical distribution area of each species using GBIF (https://www.gbif.org/) data through the rgbif package (Chamberlain and Boettiger 2017). Then, for each species we extracted BIO17 values of its distribution area with the dismo package (Hijmans, R.J, Phillips, S., Leathwick, J. and Elith, J. (2011)), which indicate precipitation values of the driest quarter, from the international database worldclim (https://www.worldclim.org/data/bioclim.html). Median values were calculated for each species. To avoid any bias on natural geographic distribution, we excluded values collected from samples in zoos, museums or laboratories. We considered a species as xeric if the median BIO17 is below 40 and mesic if the median BIO17 is over 40. For homogeneity, we also used the median of the species for the collected samples even if we have the variable for their location of capture. The biological status of the collected samples was similar whether taking the median of the species or the specific location of capture, except for *Mastomys natalensis* (Species-BIO17 is 46 and Sample-BIO17 is 0) which was only used for the sequence-based analyses due to the ambiguity of the status.

### Species selection for the subsets

For the Murinae subset, we selected Murinae species from our dataset, and further removed 3 xeric species (*Meriones unguiculatus*, *Acomys dimidiatus* and *Arvicanthus niloticus*) for equilibrating the number of mesic (6) and xeric (8) species in the dataset. This resulted in 30 samples for the expression dataset, with 5 transitions to the xeric status.

For the “within-genera” and “between-genera” subsets, we dated transitions to arid condition by using the closest relatives in the phylogeny published by (Fabre et al. 2012) and Timetree5 (Kumar et al. 2022), supplemented by specific articles for certain nodes (see rationale and references in Supplemental Fig. 1). The two datasets resulted in 32 and 31 samples, respectively, with 4 transitions to the xeric status.

### Detecting convergent changes in gene expression data

We integrated 102 RNA-seq kidney samples extracted from public repositories or produced in the lab. An automatic workflow was set up using Nextflow (version 19.04.0, April 2019) and is summarized in Supplemental Fig. 2. The scripts used to analyze the data are available here: https://gitbio.ens-lyon.fr/LBMC/cigogne/convergent_aridity_2024.

### Published RNA-seq libraries

We interrogated the NCBI for rodent Illumina RNA-seq libraries, and selected those with kidney in the metadata. To limit heterogeneity, we only selected Illumina-based RNA-seq in Genbank BioProjects. We manually removed pooled data and excluded mouse and rat data. A preliminary quality control using the top 500000 reads was performed for each sample, allowing us to select manually the three best individuals per species whenever possible (using FastQC and MultiQC (Ewels et al. 2016). The 60 selected samples, from 21 different species, are listed in Tables S1 and S4.

### *De novo* transcriptome assemblies

We generated *de novo* transcriptome assemblies for 37 species. We used the selected 69 public samples, plus 20 of our samples (Supplemental Table 3). We removed adapters and low-quality bases (Q<20) using Trimmomatic version 0.38, with options “TRAILING:20 MINLEN:25 AVGQUAL:20” (Bolger et al. 2014). After this trimming, we checked the quality of the reads with FastQC. We then assembled the data with Trinity version 2.8.5 (Grabherr et al. 2011) with option “--full_cleanup”. We predicted coding sequences from trinity assemblies with TransDecoder version 5.5.0, retaining only the best open reading frame per transcript, at least 80 amino-acids long (https://github.com/TransDecoder/TransDecoder). Basic quality values of assemblies, such as N50 and number of transcripts were retrieved with the implemented Trinity script trinityStats.pl (Haas et al. 2013). Completeness of gene repertoire was evaluated with BUSCO version 3.0.2 (Haas et al. 2013; Manni et al. 2021) with the mammalian library (mammalia_odb9). The quality of the assemblies is summarized in Supplemental Table 4.

### Quantification of expression levels

Expression levels were obtained for 34 species and 2 strains by mapping the sequence reads against coding sequences from de novo assemblies using Kallisto 0.45.1 (Bray et al. 2016) with default parameters.

### Annotation of transcripts

We selected the rodent subset from the orthology database EggNOG version 5 (Huerta-Cepas et al. 2019) and used them as a BLASTX database (containing 14 rodent genomes, that are used for annotating the families but not in our sequence and expression datasets). Coding sequences (CDSs) were aligned to this database using BLASTX (with options-outfmt ‘6 qseqid sseqid evalue bitscore length pident qstart qend sstart send’ -max_hsps 1 - max_target_seqs 1). We retained the best hit for each CDS (with an E-value threshold 1e-6), and assigned it to the corresponding EggNOG cluster. In case there were several CDS of the same species associated with a given EggNOG cluster, we retained the CDS with the best hit.

### Preparation of expression matrices

All transcripts of a given gene were imported using tximport package (Soneson et al. 2015) with option countsFromAbundance=”lengthScaledTPM” for additional scaling using the average transcript length. This accounts for gene length differences between species. We performed the following steps on each data set independently. We first adjusted the expression levels to minimize the influence of phylogenetic effect by applying batch correction using *ComBat_seq* from sva package (Zhang et al. 2020).

We defined batch groups based on species phylogenetic relatedness. A batch group usually corresponds to a rodent family. Because we need at least one xeric species and one mesic species in a batch group to perform resampling (see below), we combined two sister families in the same batch when needed (see Table S5). The correction was realized on all sets except Murinae because all species belong to the same family.

We implemented PCA using the prcomp function from the stats package, before and after batch correction. Between-class analyses were used to estimate the effect of different factors on the PCA axes (Dray and Dufour 2007).

### Convergence detection by differential and correlation network analyses

Differential expression analyses and co-expression analyses were performed on the four different data sets with their respective prepared count matrices. Only genes with non-null values in all individuals were used and mitochondrial genes were removed.

We performed differential expression analyses using the DESeq2 package (Love et al. 2014) with the following command lines:

dds <-DESeq(ddsInput)

res <-results(dds, lfcThreshold=.4, altHypothesis= “greaterAbs”)

Differentially expressed genes were filtered based on Log Fold Change threshold 0.4 corresponding to fold change over 1 (“greaterAbs”) and adjusted p-value <0.1. Thresholds were chosen after a thorough comparison (Supplemental Table 6).

We searched for modules of genes with correlated expression values with a Weighted Gene Co-expression Network Analysis (WGCNA (Horvath 2011)). We used normalized counts obtained with DESeq2 and the top 50% most variable genes based on *varianceStabilizingTransformation* from DESeq2. We selected soft-thresholding power from 16 to 20 based on the *pickSoftThreshold* function and we used the ‘signed’ network and a minModuleSize = 30 in the *blockwiseModules* function in WGCNA.

We performed functional enrichment for Gene Ontology terms and Reactome pathways using the ClusterProfiler package (Wu et al. 2021) on lists of differentially expressed genes and modules of co-expressed genes significantly correlated with xeric/mesic habitat. We also used the package REVIGO for visualization (Supek et al. 2011).

We also intersected the lists with kidney marker genes and genes involved in kidney diseases. 391 marker genes were retrieved from (Park et al. 2018; Cao et al. 2018) and manually curated from literature; 179 of these genes were available in the Murinae dataset. 244 disease genes were retrieved from the OMIM database and (Park et al. 2018), of which 165 were found in the Murinae dataset. Enrichments were computed with Fisher’s exact tests.

### Testing the significance of the number of differentially expressed genes

To estimate whether the number of observed DE genes between xeric and mesic species is significantly different from a random observation, we set up a simulation protocol inspired by (Bittner et al. 2022). This protocol preserves the overall distribution of the phenotype in the phylogeny. For each of the datasets, we defined phylogenetic groups within which the species labels can be permuted (Supplemental Table 5). The following steps were then carried out, and repeated 1000 times (See Supplementary Methods for details). Within each taxonomic group, each xeric species was associated with a mesic species randomly and individuals were subsampled to equalize the number of samples in each species of the pair. An “observed” number of differentially expressed genes was calculated with this reduced table retaining the true labels (adjusted p-value <0.1). For each gene and each previously associated pair of species, expression values were swapped between the xeric and mesic samples with a probability of 0.5. The permuted table was then used to calculate an “expected” number of DE genes. At the end of 1000 permutations, we performed a paired Wilcoxon test to compare the distribution of “observed” with those “expected” DE genes.

### Deconvolutions

To determine whether changes in cell proportions occurred between mesic and xeric species, we used computational methods to infer cell type proportions from bulk transcriptomics data. Many methodologies to infer proportions of individual cell types from bulk transcriptomics data have been developed, some of which using marker genes for different cell types, and others using scRNA-seq data. We implemented the former using sets of known marker genes plus marker genes extracted from the reanalysis of a mouse single-cell RNA-seq kidney dataset (( Park et al. 2018) and Supplemental Table 8). For the latter methods, we used the same whole scRNA-seq dataset. Upon reanalysis of these data, we removed one of the 7 individuals in the original publication. This sample (ind 7) created an additional cluster and lacked several clusters in the published parent study. Data were then normalized using SCTransform and UMAP was then generated using 15 dimensions of the PCA (Seurat package (Hao et al. 2021)). Cell type identities assigned in the original publication were then re-attributed to each cell. To determine the best deconvolution method for our data, we used the available benchmark from Cobos et al. (Avila Cobos et al. 2020) and tested 12 methods on our bulk RNA-seq data (Supplemental Fig. 9). With the best applicability to other Murinae species and good results in Mus musculus, MuSiC was selected in our analysis. Estimated proportions were then plotted per cell type and summarized by PCA analyses. Pearson correlations between estimated proportions and WGCNA eigengene values were computed and shown by heatmap.

### Detection of convergent changes in protein sequences

#### Coding sequences from genome assemblies

We retrieved coding sequences from 24 published rodent genomes (Ensembl release 99 (Harrison et al. 2024)). For nine species, kidney transcriptome assemblies and genome were available (Supplemental Table 1). In these cases, cDNA Ensembl sequences were favored over transcriptomes in the sequence analysis. We assigned these coding sequences to EggNOG groups as described above (‘Annotation of transcript’).

#### Sequence alignments

We grouped coding sequences associated with the same EggNOG group into gene families. We removed families with less than 3 species and sequences within families if their length is smaller than 70% of the median length of the family. We aligned the remaining protein sequences with MAFFT (version 7.313, with options --localpair --maxiterate 1000 (Katoh and Standley 2013)) and cleaned the alignments with HmmCleaner (version 0.180750 (Di Franco et al. 2019)). We discarded sequences for which more than 50% of the positions were removed by HmmCleaner and amino acid sites with more than 10% gaps. Finally, we back-translated the protein alignments into nucleic sequences. We obtained multiple alignments for 11437 sets of orthologs, ranging from 3 to 51 species (plus 2 strains).

#### Phylogenetic reconstruction

We selected the 4,065 complete gene families (with 53 sequences), and retained only the sites without gaps for phylogenetic analysis. Ten subsets were extracted from these families. For each subset, we chose randomly 500 genes and 200 sites per gene, and then concatenated the 100,000 sites (using catfasta2phyml.pl (https://github.com/nylander/catfasta2phyml). For genes shorter than 200 sites, all sites were retained in the concatenate. Phylogenetic reconstruction was performed using raxml-ng software (Kozlov et al. 2019) with options –all and --model LG+G. We then obtained 10 different trees. We estimated the likelihood of the complete dataset given these 10 trees and retained the tree with the best likelihood as the species tree presented Fig.1 and used it for detecting convergent sequence evolution. The chronogram presented Supplemental Fig. 1 was established with Timetree5 (Kumar et al. 2022).

#### Detection of convergent sites

We used Pelican (Duchemin et al. 2023) to detect convergent changes in protein alignments using “multinomial-filter=0.8”. Pelican annotated the tips of the species tree (see above) with xeric and mesic labels and inferred ancestral states by using parsimony. For each dataset, we filtered the gene families based on the total numbers of mesic and xeric species (see main text). We further refined the list of candidate sites by discarding all sites that had undergone a substitution in only one of the xeric clades, considering that we were interested in profile changes that have occurred in a convergent manner, at least in two xeric clades, using a custom script.

We ranked genes based on the p-value of their best site and used this ranking for gene set enrichment analyses (GSEA). GSEA were performed using Gene Ontology terms, Reactome pathways, and a custom set made of the differentially expressed genes using the ClusterProfiler package (Wu et al. 2021).

## DATA ACCESS

All raw sequencing data generated in this study have been deposited to the EBI under accession number PRJEB54931. Previously published cDNA libraries and expression raw data are listed with accession numbers in Supplemental Table 1. All codes are available in a gitlab repository (https://gitbio.ens-lyon.fr/LBMC/cigogne/convergent_aridity_2024). Sequence alignments, species tree, Pelican results, count table (used for expression analyses) and the associated coldata, as well as Supplemental Fig. 3 and 5 have been deposed on Dryad (https://doi.org/10.5061/dryad.r7sqv9sm1).

## COMPETING INTEREST STATEMENT

### Protection of Human Subjects and Animals in Research

Concerning the collection of Apodemus mystacinus in Creta, the NHMC performs collections under the provisions of the Presidential Decree 67/81. According to the Nagoya protocol, access and sharing of advantages was agreed by the government of the Republic of Benin (file 608/DGEFC/DCPRN/PF-APA/SA). The South African rodent specimens were sampled at Tussen die Riviere Nature Reserve (Free State, South Africa) in October 2017 under permit number JM 1193/2017, issued by the issued by the Free State Department of Economic, Small Business Development, Tourism and Environmental Affairs (DESTEA) in Bloemfontein (Free State, South Africa). These samples have been sent to France under export permit JM 3007/2017, also issued by DESTEA. As these species are classified as Least Concern by the IUCN, and do not require CITES permits for international transport, the samples were transferred to France under import permits issued by the Direction régionale de l’environnement, de l’aménagement et du logement (DREAL) Occitanie in Toulouse (France).

## ACKNOWLEDGMENTS

We are grateful to the following for collecting and/or giving access to their live collection: Pascale Chevret, Frédéric Delsuc, Nico Avenant and Lionel Hautier for Southern Africa samples, Gauthier Dobigny (IRD), Caroline Tatard (IRD) and Philippe Gauthier for Cameroon and Benin samples, Radim Sumbera and Lucie Plestilova for the Fukomys species, Petros Lymberakis (Natural History Museum of Crete - University of Crete) for Apodemus mystacinus, Laurent Granjon (IRD-CBGP), Youssou Niang (IRD-CBGP), Mamadou Kane, Aliou Sow, Moussa Sall, Cheikh Niang, Kodé Fall and the CERISE project for Senegal samples, François Bonhomme for ISEM collections. We also thank the GenomeEast IGBMC sequencing platform for RNAseq. This project was supported by an ANR grant to BB, MS, SP (ANR-15-CE32-0005). We thank P Veber and L Duchemin for fruitful discussions. We gratefully acknowledge support from the PSMN (Pôle Scientifique de Modélisation Numérique) of the ENS de Lyon and the French Institute of Bioinformatics (IFB CNRS UMS 3601) for the computing resources.

## Supplemental File

Supplemental file. Contains supplemental methods, lists of Supplemental Tables and supplemental figures.

